# Activity drives local CaMKII synthesis and subcellular localization via autophosphorylation-dependent pathways

**DOI:** 10.1101/2025.11.03.686364

**Authors:** Kelsey J. Clements, Nannan Chen, Kevin M De León González, Yunpeng Zhang, Avital A. Rodal, Leslie C. Griffith

**Author notes:** These authors contributed equally to this work (Kelsey J. Clements, Nannan Chen). **Corresponding Author:** Leslie C. Griffith MD PhD, Department of Biology and Volen Center for Complex Systems Brandeis University, 415 South St.m, Waltham, MA 02454-9110.

## Abstract

Strong repeated stimulation of a neuron triggers an increase in local protein synthesis along with changes in protein localization and degradation, allowing the neuron to rapidly adjust its proteome in response to activity. One of these proteins, calcium/calmodulin-dependent protein kinase II (CaMKII), is involved in mediating structural and functional changes after activity. While postsynaptic CaMKII translation has been studied, very little is known about presynaptic synthesis, including its mechanisms and functional consequences. We utilized the *Drosophila* larval neuromuscular junction (NMJ) as a model to study the molecular requirements for activity-dependent CaMKII synthesis and localization. Presynaptic-specific tagging of endogenous CaMKII demonstrates that spaced stimulation rapidly increases presynaptic CaMKII through local translation of pre-existing mRNA, independently of somatic factors. This activity-dependent synthesis requires the distal 3′ untranslated region (3′UTR) of the *CaMKII* mRNA, which is also necessary for steady-state synaptic accumulation. Additionally, we show that activity-dependent synthesis requires CaMKII T287 autophosphorylation-induced activation of the PI3K/Akt/mTor pathway. Pulse-chase experiments reveal that newly-synthesized CaMKII localizes differently from pre-existing CaMKII following stimulation, generating two overlapping CaMKII pools. We further demonstrate that a phosphomimetic T287D mutant localizes preferentially to the synaptic membrane at baseline, but cytosolic T287D protein is degraded following stimulation. Additionally, presynaptic expression of T287D-CaMKII increases spontaneous neurotransmitter release, despite lower total kinase levels. We propose a feedback model in which autophosphorylation drives both CaMKII synthesis and degradation. Synthesis enlarges the pool available for recruitment to the membrane, then degradation clears unbound activated CaMKII during rest, preserving precise control over CaMKII abundance and activity.

**Significance Statement:** Sustained synaptic changes after periods of neuronal activity require precise regulation of local protein synthesis. However, the mechanisms that control presynaptic synthesis are poorly understood. We demonstrate that spaced stimulation induces 3’UTR- and PI3K/Akt/mTor-dependent local translation of CaMKII, a key regulator of learning and memory. Additionally, CaMKII phosphorylation is involved in both synthesis and degradation of CaMKII protein. We also show that newly-synthesized and pre-existing pools of CaMKII occupy distinct regions of the bouton, suggesting that new and old CaMKII pools may have specialized functions. These findings illustrate how neuronal activity induces local protein signaling to adjust both the quantity and spatial distribution of CaMKII protein, which has lasting effects on synaptic structure and function.

## Introduction

Critical for many forms of plasticity, calcium/calmodulin-dependent protein kinase II (CaMKII) is one of the most abundant neuronal proteins in vertebrates and invertebrates (1–4). Despite its high baseline abundance, its ability to drive plastic change requires activity-dependent increases in both the absolute amount of kinase protein and its association with synapses. These changes occur in both presynaptic and postsynaptic compartments (5–7), yet detailed investigations have largely focused on dendritic local synthesis and CaMKII relocalization to the postsynaptic density in mammalian neurons. How different processes regulating dynamic changes in synaptic kinase levels interact to facilitate presynaptic plasticity is very poorly understood.

Activity-dependent relocalization couples CaMKII activation to subcellular residency, providing a mechanism to tune both the amount and location of the kinase at synapses during plasticity. Studies of cultured hippocampal neuron dendrites show that CaMKII shuttles between cytosolic, cytoskeletal, and postsynaptic pools in a manner that depends on calmodulin binding and autophosphorylation state (7–9). Activity-dependent localization to the postsynaptic density requires a conformational change that occurs when the enzyme becomes activated, and this change can be reproduced in the absence of calcium by phosphomimetic mutations of the kinase’s key regulatory autophosphorylation site T286 (T287 in *Drosophila*) (10, 11).

The underlying mechanistic assumption in studies of relocalization, which has been supported by biochemical studies of the interactome of CaMKII in different autophosphorylation states (12, 13), is that the inactive and active kinase have distinct sets of binding partners.

Catalytically dead CaMKII that is in an “active” conformation is fully capable of inducing potentiation (14), suggesting an important scaffolding function for the enzyme that is independent of its enzymatic activity. The most functionally important synaptic binding site for CaMKII in the mammalian postsynaptic density is the GluN2B C-terminus (15), but there are clearly also other dendritic ligands for active CaMKII including cytoskeletal proteins (16, 17).

CaMKII dynamics on the presynaptic side have not been extensively explored. Evidence of activity-dependent relocalization within presynaptic terminals is limited to electron microscopy studies of mammalian brain (6) and observations of clustering of a CaMKII sensor in response to calcium at the *Drosophila* larval neuromuscular junction (NMJ) (18). Given that the pre- and postsynaptic proteomes are quite different (19), the synaptic binding partners of translocated CaMKII would also be expected to be distinct in the two compartments. CaMKII has been shown to associate with synaptic vesicles (20) and in proteomic studies, while the majority of CaMKII binding partners had postsynaptic function, the defining proteins of the presynaptic active zone have also been identified as interactors (13). These studies provide potential candidate binding targets for activated presynaptic CaMKII.

In addition to redistributing CaMKII that is already present in neurons, strong or repeated activity triggers the synthesis of new CaMKII at synapses (21–24). Local translation is now recognized as a major force shaping the distinct proteomes of many subcellular compartments (25) and may even be a mechanism for minimizing energy expenditure by the cell (26). For neurons, it is thought that local translation supports plasticity and development by enabling modification of the synaptic proteome, allow it to take on new functionality. This is most easily understood for plasticity-related proteins that have very low basal levels and undergo massive upregulation with activity like Arc/Arg3.1 (27) and Homer (28). CaMKII, in contrast, is present at baseline in concentrations that are similar to those of the major cytoskeletal proteins actin (29) and tubulin (30). The need for additional protein is unclear, and whether the newly synthesized kinase participates in some preferential way in translocation to synapses is unknown.

Activity can also induce an increase in protein degradation (31) alongside protein synthesis. While autophosphorylated CaMKII has a shorter half-life than unphosphorylated CaMKII (32), it is not clear whether neuronal stimulation can specifically induce degradation of CaMKII. Protein degradation has an important role in remodeling the neuronal proteome in conjunction with synthesis, and blockade of the proteasome interferes with long-term memory formation (33). However, some models show that simultaneous inhibition of synthesis and degradation can preserve normal memory formation (34), suggesting that plasticity depends not solely on either individual process, but on the balance of protein synthesis and degradation.

In terms of mechanisms, translation of mammalian αCaMKII has been studied fairly intensively (35–38). Its mRNA was the first plasticity-related transcript shown to localize to synapses (39) and among the earliest proteins demonstrated to undergo activity-dependent local translation (40). Since then, αCaMKII translation has served as a workhorse readout for studies of local protein synthesis in mammalian dendritic processes. Very little, however, is known about presynaptic local synthesis of mammalian CaMKII, in spite of the presence of *CAMK2A* mRNA and translational machinery in that compartment (5).

In *Drosophila,* both sides of the synapse have been studied. In adult antennal lobe projection neuron dendrites, CaMKII translational reporters containing a 1260 bp proximal fragment of the 1990 bp 3’UTR demonstrated activity-regulated protein synthesis (21, 41). On the presynaptic side at the larval NMJ, activity-dependent plasticity of spontaneous release requires the distal end of the *CaMKII* 3’UTR (42). Activity at the NMJ has also been shown to increase CaMKII protein levels in presynaptic terminals, however, while the effect was blocked by cycloheximide, the authors favored a model in which synthesis was occurring in the somatic compartment of the motor neuron and protein was transported (24). In the adult mushroom body memory formation center, the axonal synaptic enrichment of CaMKII and short-term memory formation were shown to require a 23 bp translation-enhancing distal *CaMKII* 3’UTR element (43). Whether the steady-state levels of mammalian CaMKII are set up by a translational mechanism is unknown and what the relationship of activity is to steady-state synaptic enrichment is also unknown in either vertebrates or invertebrates.

In this study, we use the *Drosophila* larval NMJ to investigate these questions. We find that both steady-state and activity-dependent accumulation of CaMKII are sensitive to activity patterns and depend on the *CaMKII* distal 3’UTR. In the presynaptic terminal, strong spaced stimulation requires CaMKII autophosphorylation and an intact PI3K/mTor pathway to increase CaMKII synthesis. We also demonstrate that this new CaMKII localizes to subcellular regions distinct from those occupied by previously-synthesized CaMKII and that this combination of activation and relocalization is correlated with functional changes in transmitter release.

## Results

### Presynaptic CaMKII levels are regulated by both chronic and acute changes in activity

To investigate the mechanisms and outputs of CaMKII translational regulation we turned to the *Drosophila* larval NMJ. We first sought to determine if long-term alterations in neuronal firing could influence steady-state CaMKII protein levels by examining *Drosophila* mutants with altered excitability. The hyperexcitable *eag^1^Sh^120^*mutant had increased CaMKII antibody staining in the presynaptic region of the bouton, whereas the hypoexcitable mutant *para^ts^* had decreased CaMKII compared to wildtype (Fig. 1A-B). This suggests that activity is a required and positive driver of steady-state synaptic CaMKII levels.

**Figure 1.**
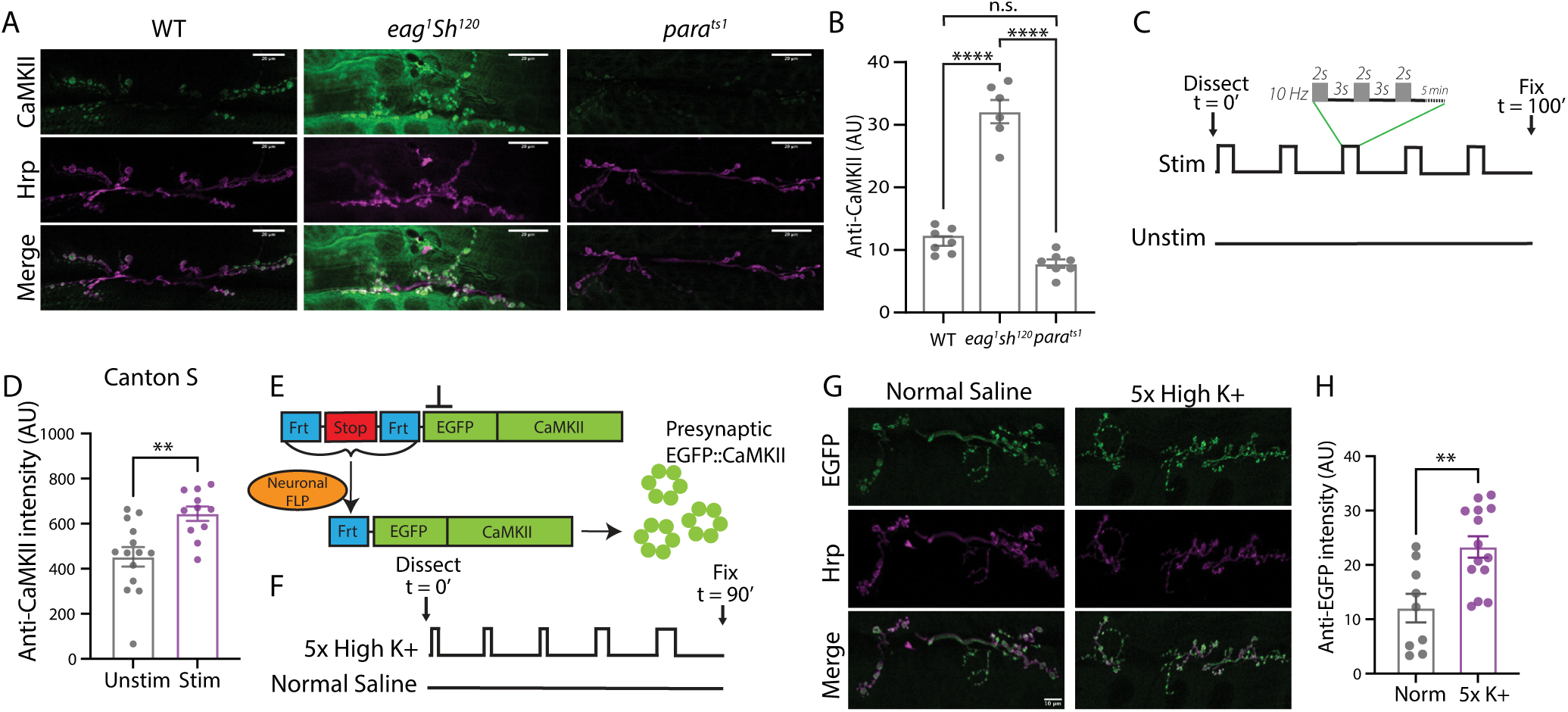
Chronic and acute activity increase presynaptic CaMKII protein at the neuromuscular junction. (A) Immunofluorescence images showing steady-state CaMKII levels in WT, *eag^1^sh^120^* and *para^ts1^*3rd instar larvae. Anti-CaMKII staining in green and Anti-Hrp staining in magenta. Scale bar = 20 µm. (B) Quantification of steady-state CaMKII levels. Differences compared with One-way ANOVA and Tukey’s multiple comparisons test, n = 7 per group. (C) Electrical stimulation paradigm. 3rd instar larvae received 5 × 5 min blocks of 10 Hz stimulation for 2 s followed by 3 s rest, repeated continuously for 5 min (40% duty cycle) with 15 min rest between blocks. (D) Quantification of anti-CaMKII staining of unstimulated or stimulated *Canton S* larvae as indicated in C, showing an increase in CaMKII protein after electrical stimulation. Differences compared with Welch’s t-test, n = 11-14. (E) Diagram of Frt-Stop-Frt-EGFP CaMKII allele, demonstrating how flies were generated that express EGFP::CaMKII solely in the presynaptic compartment. (F) 5x high potassium/spaced depolarization stimulation paradigm used to trigger CaMKII protein synthesis in larval NMJs. (G) Larvae expressing presynaptic EGFP::CaMKII were stimulated with 5x high potassium, fixed and stained with Anti-EGFP (green) and Anti-HRP (magenta) antibodies. Scale bar = 10 µm. (H) Quantification of G showing an increase in presynaptic EGFP::CaMKII after stimulation. All images are maxIP. Groups were compared with Welch’s t-test, n = 9-14. All data are reported as mean ± SEM and all P values reported as follows: * p<0.05, ** p< 0.005, *** p<0.001, **** p <0.0001.

To determine the lower limit of the timescale over which presynaptic firing can increase CaMKII levels, we electrophysiologically stimulated the motor nerve in a pattern previously shown to drive morphological plasticity at the NMJ (44). We found that delivery of 5 pulse trains to *Canton S* larvae, spaced with 15 min rest periods caused CaMKII accumulation in the presynaptic region (Fig. 1C-D). Together, these experiments implicate neuronal firing in both the acute and long-term regulation of CaMKII levels.

In the chronic and acute experiments shown above, and in the previous study examining CaMKII synthesis at the NMJ (24), colocalization of anti-CaMKII immunoreactivity with anti-HRP (a presynaptic marker) was used to assess presynaptic kinase levels. Due to the proximity of the presynaptic and postsynaptic regions, and because CaMKII is present in both compartments, it can be difficult to identify whether CaMKII staining is changing presynaptically, postsynaptically, or both with this method. To overcome this problem, we created animals containing an Frt-flanked stop element (45) followed by an in-frame N-terminal EGFP in the endogenous *CaMKII* locus (Fig. 1E). The stop cassette contains a signal which blocks transcription and translation of downstream sequences. When we express Flp recombinase exclusively in the presynaptic region, the Frt-flanked stop is removed, allowing compartment-specific expression of EGFP::CaMKII and more reliable quantification of the neuronal component of NMJ CaMKII levels.

To begin to take apart the mechanisms regulating CaMKII levels, we moved to a higher throughput method of stimulation: high K^+^ spaced depolarization (44). In this protocol, dissected third instar *Drosophila* larvae are exposed to 5 pulses of HL3 saline with elevated potassium with rest in between in normal HL3 saline (Fig. 1F). The five pulses of depolarization are analogous to the 5 stimulation trains that we delivered with an electrode in Fig. 1C. Importantly, the functional plasticity outputs of chemical and electrical/optogenetic spaced stimulation have been shown to be the same at this synapse (44). This temporal pattern of stimulation invokes the same signaling pathways that are used to form protein synthesis-dependent long-term memory: equivalent stimulation or training given in a single session produces primarily short-term, non-protein synthesis-dependent plasticity (46, 47). Applying the spaced protocol, we found that presynaptic EGFP::CaMKII increased in stimulated NMJs (Fig. 1G-H), indicating a fast activity-dependent effect on neuronal kinase levels, consistent with previous reports using antibodies (24).

### Calcium-dependent local protein synthesis drives acute increases in CaMKII

To determine if the acute effect was calcium-dependent, we asked if we could block CaMKII accumulation when calcium ions were chelated by including EGTA in the normal and stimulation solutions (Fig. 2A). We found that lack of extracellular calcium blocked CaMKII increases (Fig 2C, indicating that depolarization-induced intracellular calcium signaling is likely involved in CaMKII upregulation.

**Figure 2.**
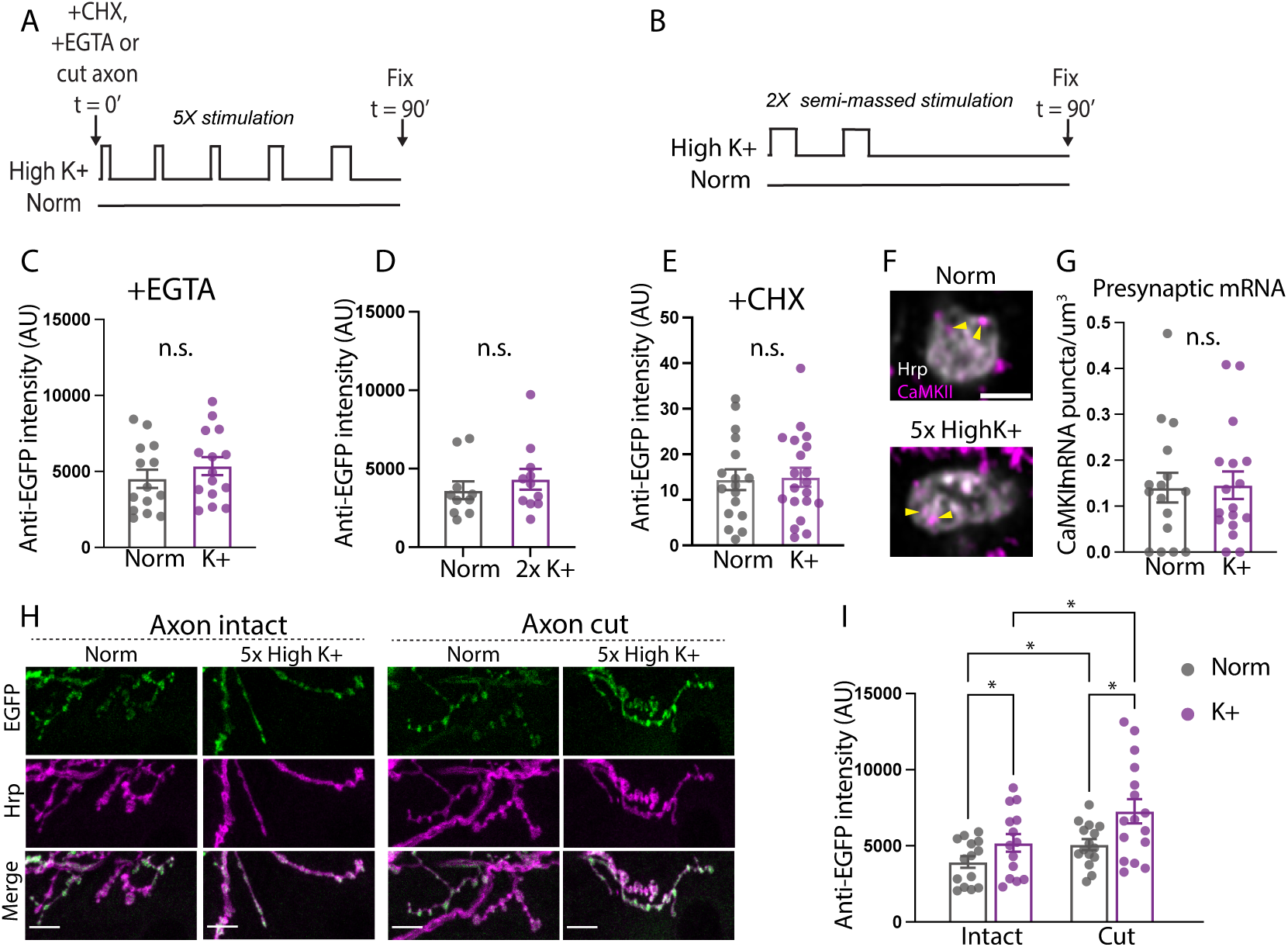
Calcium-dependent local protein synthesis drives spaced stimulation-induced increase in presynaptic CaMKII. (A) Diagram demonstrating the timing of drug application. (B) Diagram showing 2x high potassium stimulation protocol. (C, E, G) EGFP::CaMKII larvae were treated with EGTA (C), CHX (E), or stimulated with 2x high potassium (D), then fixed and stained with Anti-EGFP and Anti-Hrp antibodies to determine presynaptic EGFP::CaMKII quantity. Quantification of NMJs show lack of CaMKII synthesis after exposure to EGTA, 2x K+ stimulation or CHX. Groups were compared with Welch’s t-test, n = 14-15 for C, n = 10 for D, n = 16-20 for E. (F) Larvae were stimulated with 5x high potassium protocol, fixed, and *CaMKII* mRNA (magenta) detected with HCR-FISH, and then co-stained with Anti-Hrp antibody (grey). Scale bar = 2 µm. (G) Quantification of the number of mRNA puncta per µm^3^. There was no change in mRNA quantity due to stimulation. Data were compared with Welch’s t-test, n = 16-17. (H) Larvae were dissected, axons cut on one side, stimulated, and then stained for EGFP (green) or Hrp (magenta). Scale bar = 10 µm. (I) Quantification of Anti-EGFP intensity of NMJs represented in panel K showing normal CaMKII synthesis occurs regardless of axotomy. All images are MaxIP except in F, which are single Z slices. Significance calculated with a 2-way ANOVA and Tukey’s multiple comparisons test, n = 14-16. All data are reported as mean ± SEM and all P values reported as follows: n.s. = not significant, * p<0.05, ** p< 0.005, *** p<0.001, **** p <0.0001.

To ask whether the timing of the stimulation pulses mattered, we tested a semi-massed protocol, in which the total amount of high K^+^ stimulation time was the same as the spaced protocol (16 min), but was separated into two pulses instead of five (Fig. 2B). We observed no increase in presynaptic CaMKII after the 2x semi-massed protocol (Fig. 2D), indicating that spaced, repetitive stimulation is required, similar to other types of long-lasting plasticity that are dependent on protein synthesis (42, 44, 48).

There were several possible mechanisms that could explain the increase in CaMKII after calcium influx, including protein transport from the soma (the model favored in 24), mRNA transport to the synapse followed by local translation or local translation of mRNA already present at the synapse. To distinguish these possibilities, we first applied cycloheximide (CHX), a translation inhibitor, during stimulation. We observed no increase in CaMKII (Fig. 2E), indicating that activity-dependent CaMKII accumulation is not dependent on some pool of pre-synthesized protein, but rather requires new translation.

To determine if there was *CaMKII* mRNA present at the synapse and whether its levels changed with stimulation, we used Hybridization Chain Reaction Fluorescent In Situ Hybridization (HCR-FISH) with a *CaMKII*-specific probe. *CaMKII* mRNA could be visualized in the resting presynaptic motor terminal, and there was no increase in *CaMKII* mRNA content after stimulation (Fig. 2F-G). This argues that new CaMKII is likely coming from translation of previously localized mRNA. To confirm that there was no other somatic contribution to CaMKII synthesis, we detached the cell body from the synaptic compartment by severing the axon before stimulation. While we observed an increase in CaMKII in unstimulated axotomized NMJs possibly due to damage to the axon, we also observed a post-stimulation increase in presynaptic CaMKII. This provides evidence that mRNA and translational machinery components that are already present at the synapse before stimulation drive increases in CaMKII protein (Fig. 2H).

### The distal 3’UTR of the *CaMKII* mRNA is required for steady-state and long-term activity-dependent CaMKII synthesis

*CaMKII* mRNA has two different 3’UTRs that are produced by alternative polyadenylation: a short isoform (123 bp) and a long isoform (1990 bp). Previous work in our lab demonstrated that only the long 3’UTR form of *CaMKII* mRNA, but not the short 3’UTR form, can support synaptic enrichment of CaMKII protein in the *Drosophila* mushroom body (43). To ask if steady-state accumulation of CaMKII at the NMJ also requires the 3’UTR, we utilized flies from that study in which the wildtype 3’UTR was either deleted (*CaMKII^UDel^*) or reduced to the short isoform (*CaMKII^UShort^*) by mutations made in the endogenous locus. As a genetic control, we used a line (*CaMKII^ULong^*) in which the full long 3’UTR was restored on the *CaMKII^UDel^* background. Compared to either *Canton S* wild type or the control *CaMKII^ULong^*, *CaMKII^UDel^* and *CaMKII^UShort^* larvae had significantly reduced steady-state levels of CaMKII (Fig. 3A-B). Because the stability of the mRNA is not affected by these UTR deletions (43), this result argues that the steady-state accumulation of CaMKII at NMJ may be controlled by translational regulation.

**Figure 3.**
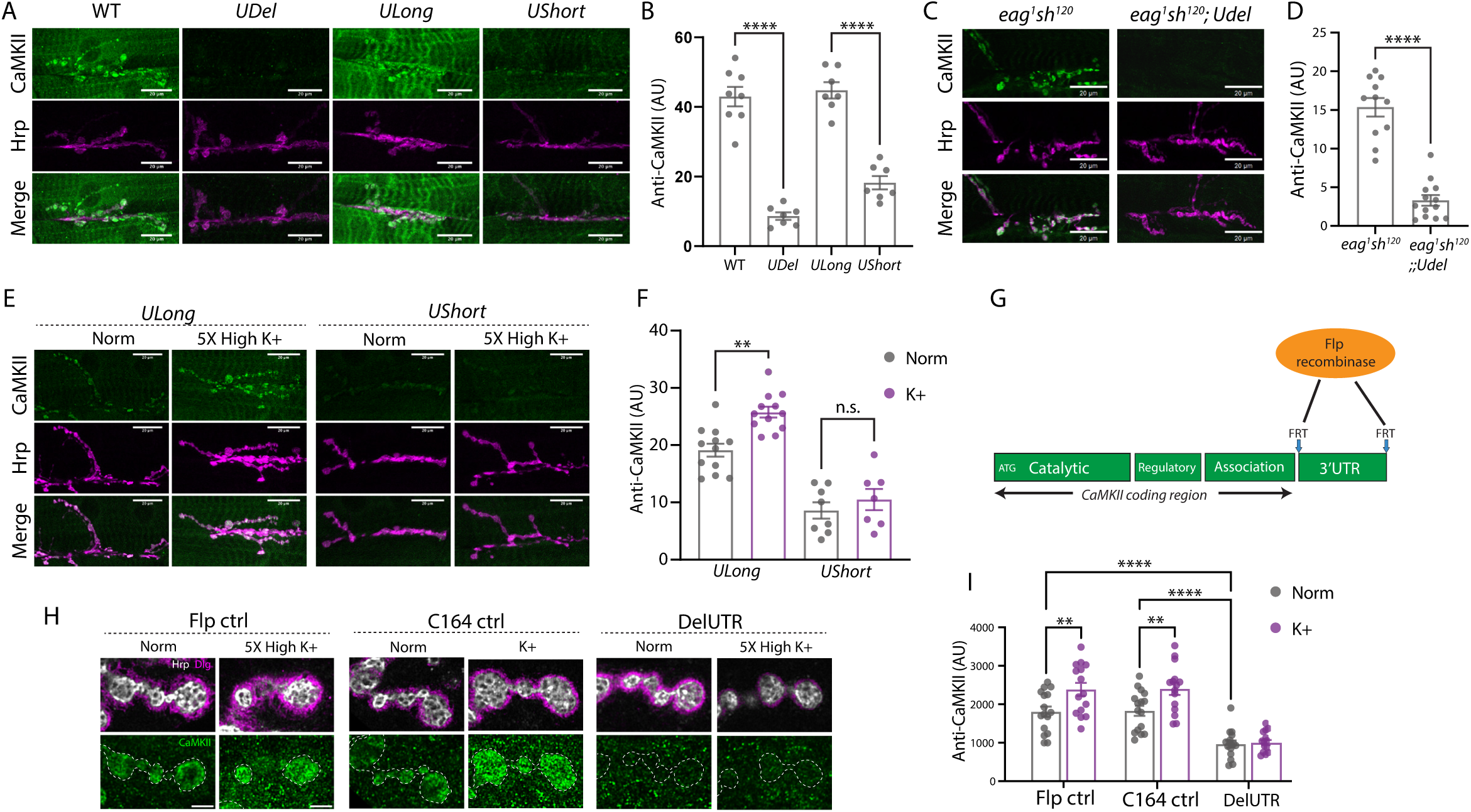
The Long form of the *CaMKII* 3’UTR is required for steady-state and activity-dependent CaMKII synthesis. (A) WT, *CaMKII^Udel^*, *CaMKII^UShort^*or wild type *CaMKII^ULong^* larvae were dissected, fixed and stained with Anti-CaMKII (green) and Anti-HRP (magenta) antibodies. (B) Quantification of fluorescence intensities of NMJs represented in A. One-way ANOVA with Tukey’s multiple comparisons test, n = 7-8. (C) *eag^1^sh^120^*or *eag^1^sh^120^*; *CaMKII^UDel^* larvae were dissected, fixed and stained with Anti-CaMKII (green) and Anti-HRP (magenta) antibodies. (D) Quantification of larvae in C, showing the UTR is required for steady state CaMKII expression. Groups compared with an unpaired t-test, n = 11-13. (E) *CaMKII^UShort^* or wild type *CaMKII^ULong^* larvae were dissected and exposed to 5x high potassium protocol, then fixed and stained with Anti-CaMKII (green) and Anti-HRP (magenta) antibodies. (F) Quantification of larvae represented in panel E. Groups compared with a One-way ANOVA with Tukey’s multiple comparisons test, n = 8-12. Scale bars = 20 μm. (G) Schematic demonstrating how CaMKII-Frt-3’UTR-Frt flies were used to delete the 3’UTR in the presynaptic region. (H) Representative images of high resolution Airyscan imaging of Flp or C164 control, or DelUTR NMJs that were stimulated, fixed and stained with Anti-CaMKII (green), Anti-HRP (grey), or Anti-Dlg (magenta) antibodies. Presynaptic regions are outlined. Scale bar = 3 μm. (I) Quantification of fluorescence intensities from experiment described in panel H, showing along with panel F, that the long UTR is required for activity-dependent expression. All images are MaxIP except 3H which are single Z slices. Data were compared with a 2-way ANOVA and Tukey’s multiple comparisons test, n = 15-16. All data are reported as mean ± SEM and all P values reported as follows: n.s. = not significant, * p<0.05, ** p< 0.005, *** p<0.001, **** p <0.0001.

We next wanted to look at long-term effects of activity on CaMKII in animals with 3’UTR deletions. The increased expression of CaMKII observed in *eag^1^Sh^120^* larvae was blocked in *eag^1^Sh^120^*;;;*CaMKII^UDel^*animals (Fig. 3C-D) and spaced depolarization-stimulated increases were not seen in *CaMKII^UShort^*(Fig. 3E-F). This dependence on the distal *CaMKII* 3’UTR supports the idea that translation in neurons at the NMJ is a critical regulator of activity-dependent and steady-state CaMKII levels.

While the findings with *CaMKII* UTR mutants demonstrate that the distal part of the 3’UTR is involved in activity-dependent synthesis as well as steady-state synaptic enrichment, these data do not clearly separate presynaptic from postsynaptic mechanisms since the 3’UTR was mutated in the entire animal. To determine whether the 3’UTR on the presynaptic side is required for the activity-dependent synthesis of CaMKII, we deleted the entire 3’UTR selectively in motor neurons using a line in which Frt sites were inserted at the beginning and end of the 3’UTR (Fig. 3G). Flp recombinase was expressed using a presynaptic driver (*C164-Gal4*), to delete the 3’UTR only in the motor neurons while leaving the postsynaptic muscle 3’UTR intact. After spaced depolarization, stimulated genetic control larvae demonstrated higher levels of CaMKII compared to unstimulated larvae, but there was no increase in CaMKII protein in stimulated larvae with a deleted presynaptic 3’UTR (DelUTR) (Fig. 3H-I). Additionally, unstimulated DelUTR larvae had decreased steady-state amounts of CaMKII protein compared to genetic controls, demonstrating that the UTR is required cell autonomously for both activity-dependent and steady-state synthesis of CaMKII in presynaptic terminals.

### Acute activity-dependent CaMKII synthesis requires autophosphorylation of CaMKII

We next sought to determine the link between calcium influx into the presynaptic terminal and stimulation of CaMKII synthesis. One obvious candidate mediator is CaMKII itself. CaMKII has been shown to stimulate protein synthesis in neurons through interactions with cytoplasmic polyadenylation element-binding protein (CPEB) (49), eIF4E (50) and the mTOR pathway (51, 52). Additionally, autophosphorylation of CaMKII at threonine T287, which contributes to formation of long-term plasticity (53), can be amplified by repetitive stimulation (54). It was therefore possible that CaMKII activity is involved in the spaced depolarization-induced synthesis of its own protein at the NMJ. To explore this, we used flies in which the phosphorylation at threonine 287 in the regulatory domain of CaMKII is blocked by mutation to alanine (*CaMKII^T287A^*). The T287A mutant kinase is able to respond normally to stimulation by calcium to phosphorylate substrates, but cannot become calcium-independent. We found no high K^+^-induced CaMKII synthesis in two independently-derived *CaMKII^T287A^* lines, one of which has all three regulatory domain phosphorylation sites blocked (*CaMKII^T287AT306/7A^*) (Fig. 4A-C). This indicates that generation of calcium-independent CaMKII by phosphorylation of T287 is required for normal activity-dependent CaMKII synthesis.

**Figure 4.**
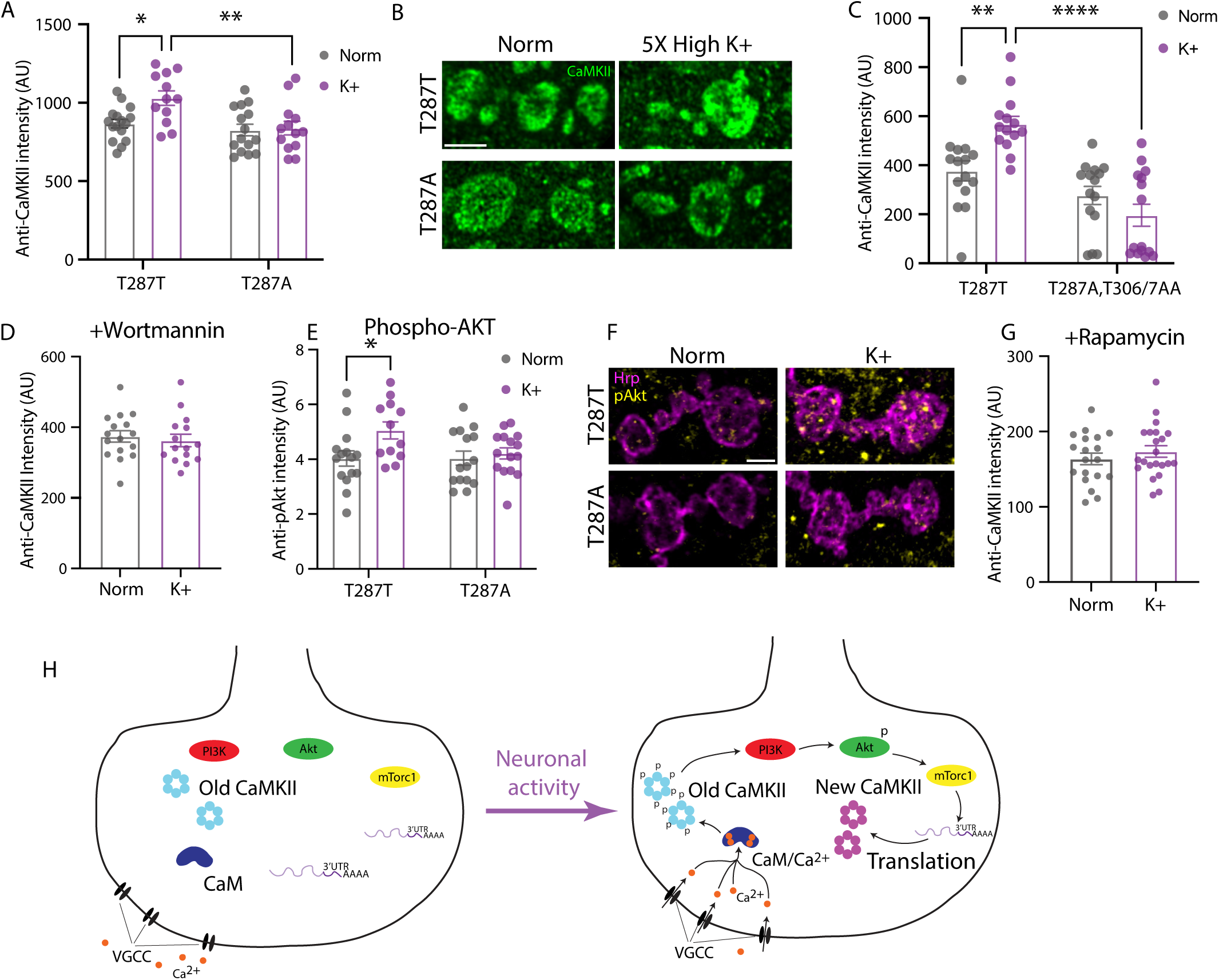
CaMKII autophosphorylation acts through the Pi3K/Akt/mTor pathway to regulate CaMKII synthesis. (A) CaMKII synthesis is blocked when the T287 site cannot be phosphorylated. *CaMKII^T287T^* wild type control or *CaMKII^T287A^* larvae were dissected, stimulated with 5x high potassium, fixed and stained with Anti-CaMKII (green). The data show Groups were compared with a 2-way ANOVA and Tukey’s multiple comparisons test, n = 12-16. (B) Representative images of NMJs from panel A. Scale bar = 3 μm. (C) Repeated experiment as in panel A except *CaMKII^T287A^* larvae were replaced with triple mutant *CaMKII^T287A,T306/7A^* larvae, n = 14-15. (D, G) CaMKII synthesis is blocked with PI3K or mTor inhibition. Larvae were pre-treated with Wortmannin (D) or Rapamycin (G) for 15 min, then stimulated with 5x high potassium, with the drug present in all solutions. Larvae were processed as in previous experiments and the anti-CaMKII signal quantified. Groups were compared using a Welch’s t-test, n = 15-16 for D and n = 19-22 for F. (E) Same experimental setup as described in panel A, except larvae were stained with a phospho-Akt antibody (yellow), demonstrating that Akt phosphorylation requires CaMKII T287 phosphorylation. Groups were compared with a 2-way ANOVA and Tukey’s multiple comparisons test, n = 12-15. (F) Representative images from panel E. Scale bar = 2 μm. All images are single Z slices. (H) Proposed model for presynaptic activity-dependent synthesis. Before stimulation, the components required for activity-dependent synthesis are present in the bouton. During stimulation, calcium ions enter the cell and bind to calmodulin (CaM), which can activate pre-existing CaMKII, leading to autophosphorylation at T287. Phosphorylated CaMKII activates the PI3K/Akt/mTor pathway, leading to 3’UTR-dependent translation of *CaMKII* mRNA. The new CaMKII protein localizes to a different region of the bouton than the old protein. All data are reported as mean ± SEM and all P values reported as follows: n.s. = not significant, * p<0.05, ** p< 0.005, *** p<0.001, **** p <0.0001.

### Autophosphorylated CaMKII controls protein synthesis via the PI3K pathway

The constitutively active autophosphorylated form of CaMKII had previously been shown to activate PI3K signaling at the *Drosophila* NMJ and increase synaptic growth (55, 56). Since PI3K can increase protein synthesis through mTor activation, this could provide an explanation for why blocking CaMKII T287 phosphorylation inhibited protein synthesis. To test the involvement of this pathway, we treated wildtype larvae with Wortmannin, an inhibitor of PI3K, during spaced depolarization, and observed no increase in CaMKII in stimulated larvae (Fig. 4D), implicating this signaling system in activity-dependent local protein synthesis. Activation of mTor-dependent translation by PI3K occurs via its ability to activate the protein kinase Akt. Consistent with this pathway being involved in CaMKII local synthesis, we found that spaced stimulation increased phosphorylation of Akt at its activation site in control larvae, but this pAkt increase was absent in *CaMKII^T287A^* larvae (Fig. 4E-F). In congruence with the previous two results, application of rapamycin, an inhibitor of mTor, the downstream link between PI3K and translation, also blocked the ability of spaced stimulation to increase CaMKII levels (Fig. 4G).

These findings support a model (Fig. 4H) in which depolarization-induced phosphorylation of CaMKII activates the PI3K/Akt/mTor pathway to stimulate CaMKII synthesis in presynaptic neurons.

### Newly synthesized CaMKII localizes closer to the active zone

CaMKII is an abundant protein and it is not clear why long-term plasticity requires synthesis of additional enzyme. We wondered if differences in phosphorylation state or interacting partners, could lead newly synthesized kinase (“new CaMKII”) to have a distinct localization compared to CaMKII synthesized before the activity pulse (“old CaMKII”). To detect possible differences in the localization of new and old CaMKII, we devised a pulse-chase method using sequential applications of different HaloTag ligands. This type of method has previously been used to measure protein synthesis, turnover and trafficking in the mammalian brain (57–59).

We expressed Halo::CaMKII presynaptically from the endogenous locus using the Flp-Frt system (Fig. 5A). After dissecting 3rd instar larvae, we applied the pulse ligand (JF646), which stains all the cell’s CaMKII. After washing, we applied the chase ligand (JF549) at a 4X concentration (Fig. 5B). This higher concentration loads the presynaptic cell with the new ligand and ensures it outcompetes any residual pulse ligand, resulting in selective labelling of newly synthesized Halo::CaMKII with the second color. We then conducted spaced depolarization to induce new synthesis, fixed, and used high resolution Airyscan imaging to visualize the two pools of CaMKII. We observed an increase in new Halo::CaMKII in the K+ stimulated condition but not in the normal control condition. Application of either cycloheximide (CHX, Fig. 5C) or anisomycin (ANISO, Fig. 5D) blocked the appearance of the new CaMKII, indicating that this increase in new Halo::CaMKII is likely due to new protein synthesis.

**Figure 5.**
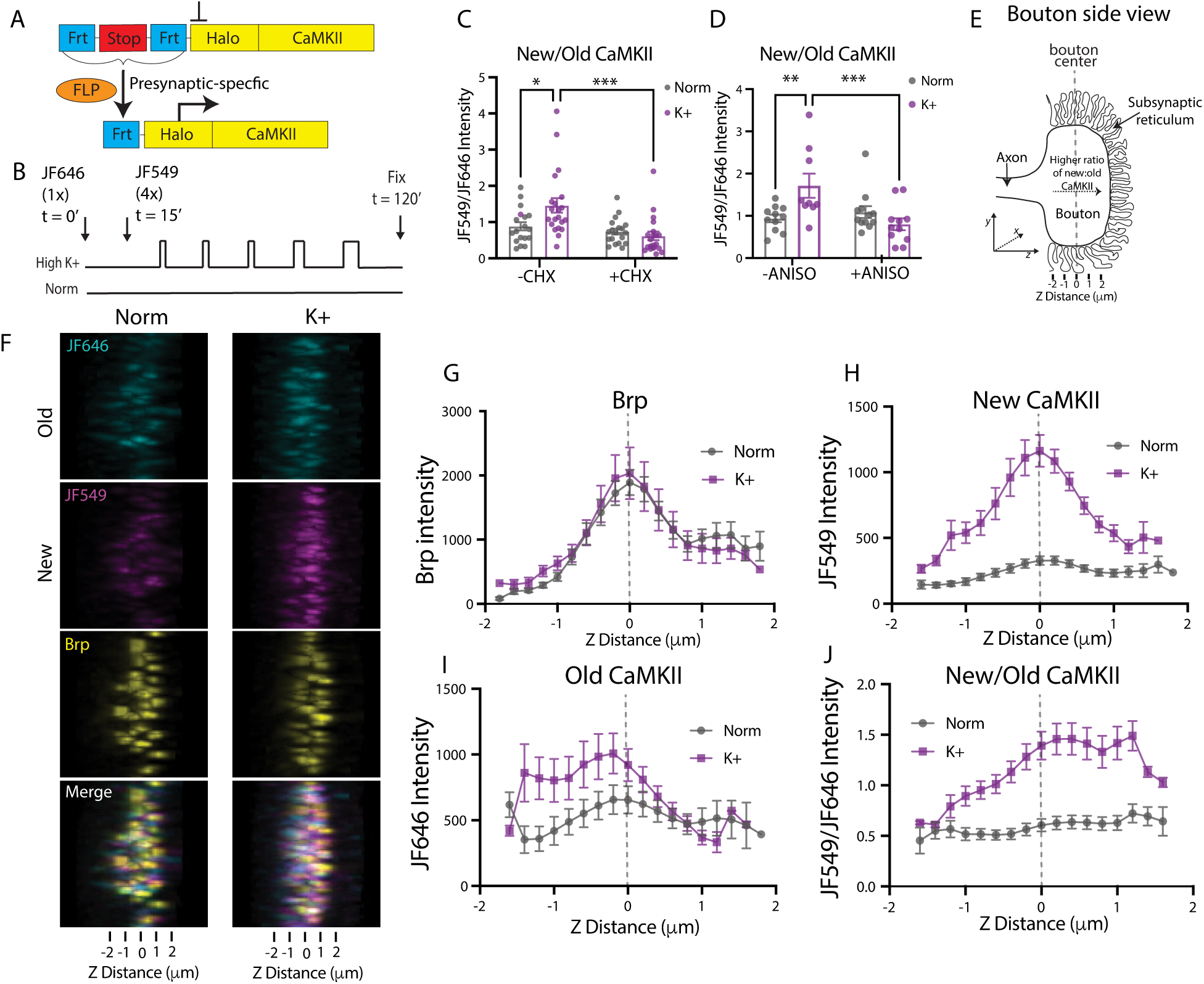
Newly-synthesized CaMKII localizes to a distinct region of the presynaptic bouton relative to pre-existing CaMKII. (A) Diagram showing strategy for conditional labelling of the presynaptic pool of CaMKII using a Frt-Stop-Frt-Halo-CaMKII cassette in the endogenous *CaMKII* locus. (B) Stimulation paradigm used for pulse chase experiments. Larvae were dissected, loaded with pulse and chase HaloTag ligands, stimulated with 5x high potassium and then fixed. (C, D) Protein synthesis inhibitors block increase in new/old CaMKII ratio. Larvae were treated as in B, and half were exposed to cycloheximide (C), or anisomycin (D) for the duration of the experiment. The ratio of new/old was calculated for each condition. Protein synthesis inhibitors block activity-dependent accumulation of new CaMKII. Differences between groups were compared using 2-way ANOVA and Tukey’s multiple comparisons test, n = 18-22 for C and 9-12 for D. P values reported as follows: n.s. = not significant, * p<0.05, ** p< 0.005, *** p<0.001, **** p <0.0001. (E) Schematic illustrating the presynaptic bouton’s location relative to the subsynaptic reticulum viewed along the Z axis. (F) Representative images from larvae stimulated and stained as described in Fig. 5A-B. View of boutons was rotated so that the Z axis can be seen. Cyan signal marks the pre-existing (old) CaMKII, magenta signal marks the newly-synthesized (new) CaMKII and yellow is anti-BRP to mark active zones. (G-J) Quantification of BRP (G), new (H), old (I) and new/old (J) CaMKII intensity across the Z axis. All data are reported as mean ± SEM.

To compare the localization of newly synthesized and preexisting Halo::CaMKII, we first examined their fluorescence distributions along the Z-axis within individual boutons. This analysis allowed us to ask whether either pool was preferentially localized toward the axon or the postsynaptic subsynaptic reticulum. We plotted the Z-axis profile of Brp staining, of which the peak was used to determine the central Z plane of the bouton (Fig. 5E-G). We then aligned the Z-stacks of all the boutons so that we could plot the Z-axis profile of each fluorophore and compare the distribution of tagged CaMKII molecules. This analysis indicated that new CaMKII of K+-stimulated boutons was distributed symmetrically throughout the bouton, with the intensity peak located at the center of the bouton, corresponding to the peak of Brp staining (Fig. 5G-H).

Intriguingly, the old CaMKII molecules in stimulated boutons became more asymmetrically distributed (Fig. 5I), shifting toward the cytosolic side of the bouton where it buds from the axon in K+-stimulated animals. In controls treated with normal saline, old CaMKII remained symmetrically distributed. The area under the fluorescence distribution curve did not differ significantly between K⁺-stimulated and control boutons (Fig. S1), suggesting that activity alters the distribution of preexisting CaMKII, resulting in greater enrichment on the axon-facing side of the bouton but no detectable net change in its abundance. This asymmetric distribution results in a higher ratio of new to old CaMKII near the segment of the bouton facing the postsynaptic subsynaptic reticulum, and a lower ratio near the cytosolic portion of the bouton (Fig. 5J). The distinct localizations of newly-synthesized and pre-existing CaMKII subunits indicate that they may form two pools of protein with differences in phosphorylation and/or binding partners that are regulated independently by neuronal activity.

### Autophosphorylation of CaMKII alters its subcellular localization

Activation of CaMKII results in a rapid autophosphorylation of a threonine (T286 in mammals, T287 in *Drosophila*) that allows the kinase to remain in an ‘open’ conformation and maintain substrate phosphorylation even after dissociation of calmodulin (11, 60–62). To determine if the spaced depolarization paradigm increased net pT287, we employed a phospho-specific antibody and observed an increase in autophosphorylated total CaMKII in stimulated NMJs (Fig. 6A-B).

**Figure 6.**
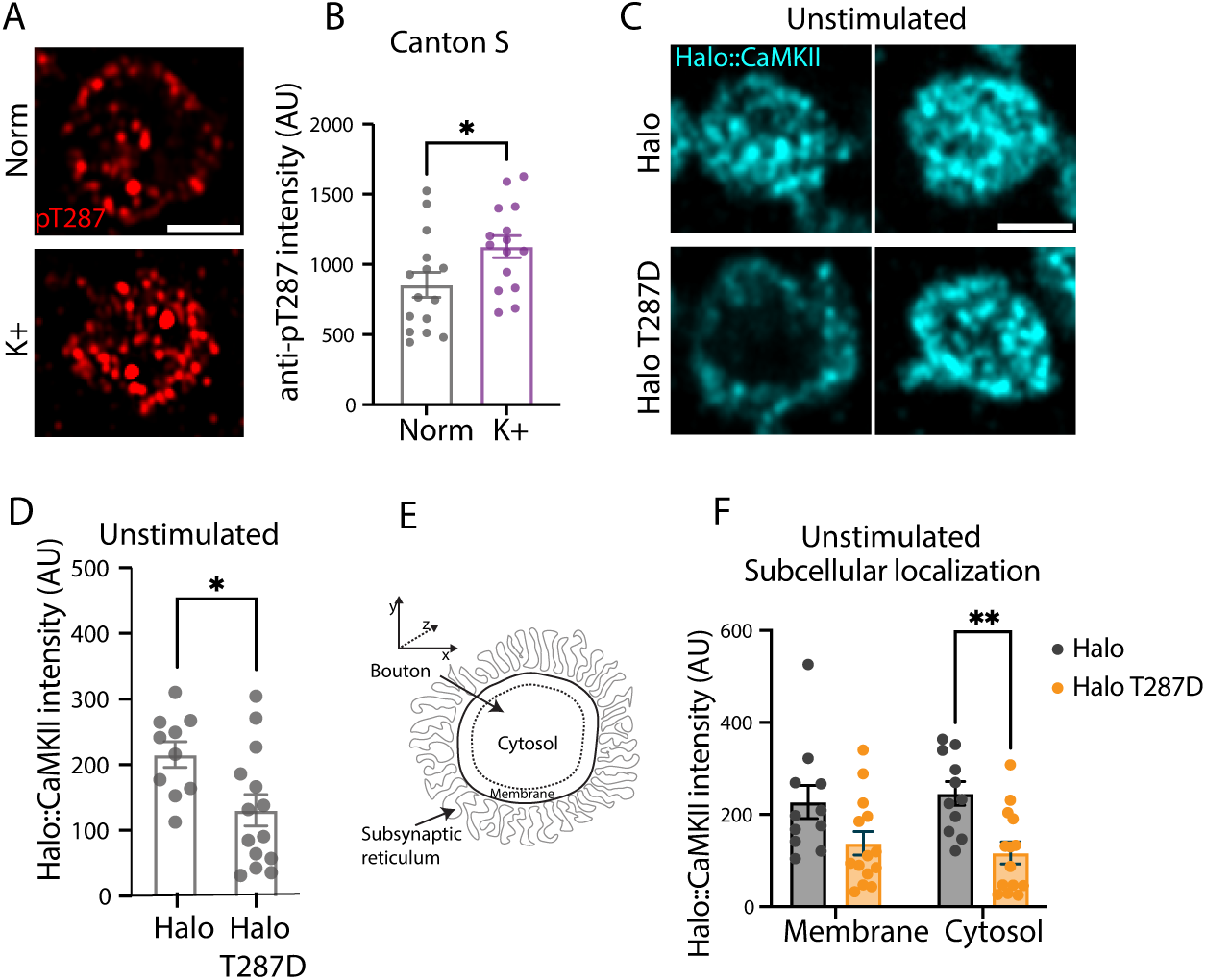
Halo::CaMKII^T287D^ is less stable and localizes to the synaptic membrane. (A) Representative images of *Canton S* NMJs stained with anti-pT287 antibodies (red) in normal control and K+ stimulated larvae. Scale bar = 2 μm. (B) Quantification of NMJs represented in panel A. Groups were compared with a Welch’s t-test, n = 15, 15. (C) Representative images Halo::CaMKII and Halo::CaMKII^T287D^ unstimulated boutons stained with HaloTag ligand JF549 (cyan). Two images for each group are included to display the variability in localization. (D) Quantification of HaloTag intensity of Halo::CaMKII and Halo::CaMKII^T287D^ unstimulated boutons. Groups were compared with a Welch’s t-test, n = 10, 14. (E) Diagram of a bouton showing the locations of the membrane and cytosolic regions compared to the postsynaptic subsynaptic reticulum for analysis in the X-Y plane. (F) Quantification of the Halo::CaMKII intensity within the membrane and cytosolic regions. All images are single Z slices. Scale bar = 2 μm. Groups were compared with a 2-way ANOVA with Tukey’s multiple comparisons test, n=10 for Halo, n=14 for Halo-T287D. All data are reported as mean ± SEM and all P values reported as follows: n.s. = not significant, * p<0.05, ** p< 0.005, *** p<0.001, **** p <0.0001.

Autophosphorylated CaMKII has been shown to dynamically relocate in response to activation in many systems, including the fly (18). In mammalian excitatory neurons, activated autophosphorylated CaMKII translocates to the postsynaptic density, where it can interact with substrates and act as a scaffolding protein (7, 63, 64). Likewise, in the presynaptic region of hippocampus, a burst of activity can cause redistribution of CaMKII, positioning it closer to synaptic vesicles and the active zone (6). To determine if autophosphorylation could alter steady-state subcellular localization of CaMKII, we generated a T287D phosphomimetic Halo::CaMKII allele which can be conditionally expressed from the endogenous locus in the presynaptic cell (as in Fig. 5A, but with the T287D mutation instead of wildtype CaMKII). Mutation of T287 to a aspartate (T287D) mimics autophosphorylation, making the enzyme constitutively active (11).

In unstimulated neurons, total Halo::CaMKII^T287D^ protein was present at lower levels than wild type (Fig.6C-D), consistent with our recent finding that adoption of an activated conformation stimulates degradation of the CaMKII holoenzyme (32). We hypothesized that interactions with different binding partners could partition CaMKII into membrane-associated and cytosolic pools. To visualize this, we needed to utilize a slightly different method of analysis from that in Figure 5 and image in the X-Y plane. We quantified Halo::CaMKII and Halo::CaMKII^T287D^ in two distinct regions of unstimulated neurons: 1) near the bouton membrane, where active zones and membrane-associated CaMKII-binding protein are located, and 2) in the bouton cytosol, 250-300 nm away from the synaptic membrane (Fig. 6E). We found Halo::CaMKII^T287D^ was relatively more concentrated at the synaptic membrane (Fig. 6F and Fig. 7C), while the wild type Halo::CaMKII protein was more concentrated in bouton cytosol (Fig. 7A-B). This may suggest that degradation of activated CaMKII is occurring primarily in cytosol.

**Figure 7.**
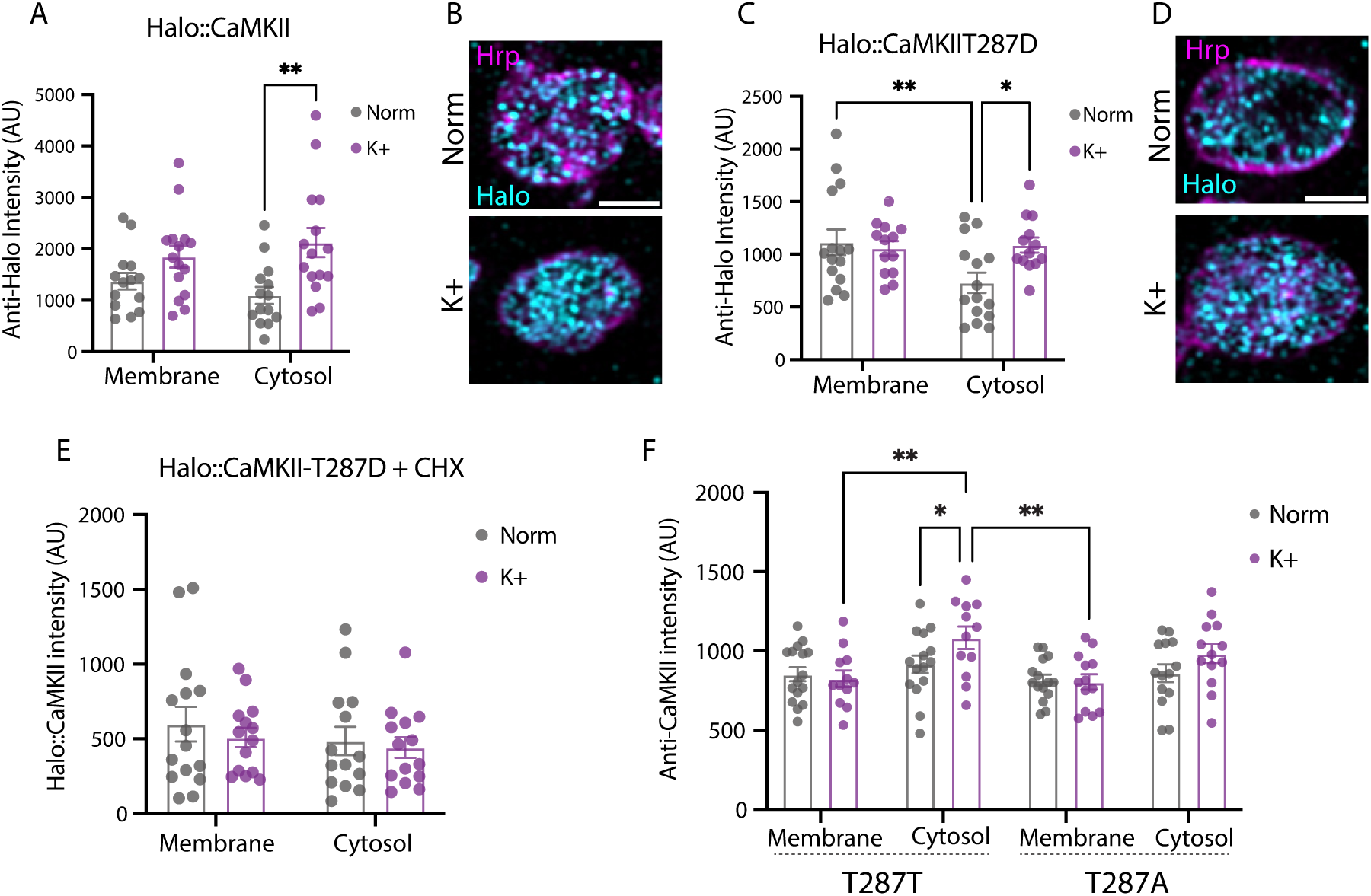
Synthesis of Halo::CaMKII and Halo::CaMKII^T287D^ occurs in the cytosolic portion of boutons. (A-D) Halo::CaMKII and Halo::CaMKII^T287D^ were stimulated and quantified as in Fig. 6E-F. (A) Quantification of Halo ligand intensity within the membrane and cytosolic regions for Halo::CaMKII (A) or Halo::CaMKII^T287D^ (C). n = 14 Norm Halo, n = 15 Halo K+, n =15 Norm T287D, n= 13 K+ T287D. (B-D) Representative images of single boutons from panels A and C. Images are single Z slices, scale bars = 2 μm. (E-F) Increase in cytosolic CaMKII is synthesis-dependent. (E) Inhibition of translation of Halo::CaMKII^T287D^ NMJs show no increase in cytosolic Halo staining. n=14,14. (F) Activity-dependent increase in cytosolic CaMKII is observed in control T287T NMJs, but not synthesis-deficient T287A NMJs. n= 16 Norm T287T, n= 13 K+ T287T, n= 15 Norm T287A, n= 13 K+ T287A. Groups were compared with 2Way ANOVA with Tukey’s multiple comparisons test. Data are reported as mean ± SEM and all P values reported as follows: n.s. = not significant, * p<0.05, ** p< 0.005, *** p<0.001, **** p <0.0001.

### Activity alters subcellular pools of CaMKII via new protein synthesis

Our data indicate that CaMKII autophosphorylation is required for synthesis of its own protein (Fig. 4), and that the phosphomimetic T287D form of CaMKII is present at lower steady-state levels compared to wildtype (Fig. 6C-D). If activity-induced T287 phosphorylation is able to induce both synthesis and degradation of CaMKII, what is the net effect of autophosphorylation on protein quantity and localization? We used HaloTagged CaMKII to track the localization of wildtype and T287D protein after stimulation to try to disentangle these competing processes. Spaced depolarization increased total (old + new) wild type Halo::CaMKII protein primarily in the cytosolic region (Fig. 7A-B), with no change in the membrane bound fraction. This pattern was also observed in *Canton S* NMJs stained for total CaMKII (Fig. S2C-D) or pT287 antibodies (Fig. S2A-B), indicating our Halo::CaMKII allele is not disrupting basic processes.

Similar to wild type, cytosolic levels of Halo::CaMKII^T287D^ increased, while membrane protein, which was higher than cytosol in the unstimulated control, remained the same (Fig. 7C-D). We hypothesized that the site of synthesis of CaMKII may reside in the cytosolic portion of the bouton. To rule out the possibility that the increases were due to a dynamic relocation of CaMKII to the cytosol in response to stimulation we applied a translation inhibitor, CHX. This blocked any increase in CaMKII in the membrane or cytosol (Fig. 7E), indicating the cytoplasmic changes are synthesis-dependent, not purely stimulation-dependent, changes. We also examined the localization of total CaMKII protein in *CaMKII^T287A^* NMJs, which are deficient in activity-dependent CaMKII translation, and in the *CaMKII^T287T^* wild type control.

While cytosolic, but not membranous, CaMKII increased in stimulated *CaMKII^T287T^* NMJs, the accumulation of cytosolic CaMKII was absent in *CaMKII^T287A^*stimulated NMJs (Fig. 7F), confirming the cytosolic increase requires new synthesis.

### Phosphomimetic CaMKII synthesized in the bouton cytosol is degraded after repetitive stimulation

Since the steady-state (Fig. 6C) and post-stimulation (Fig. 7) localization of Halo::CaMKII^T287D^ differ, we next asked whether we could visualize a post-stimulation reorganization of Halo::CaMKII^T287D^ with a longer period rest after stimulation. We quantified the localization of Halo::CaMKII^T287D^ at either our standard 15 minutes or 1 hour after the final potassium pulse (Fig. 8A) and found that while there was a trend toward decreased membranous protein (p = 0.09), there was a substantial significant decrease in the cytosolic protein (Fig. 8C).

**Figure 8.**
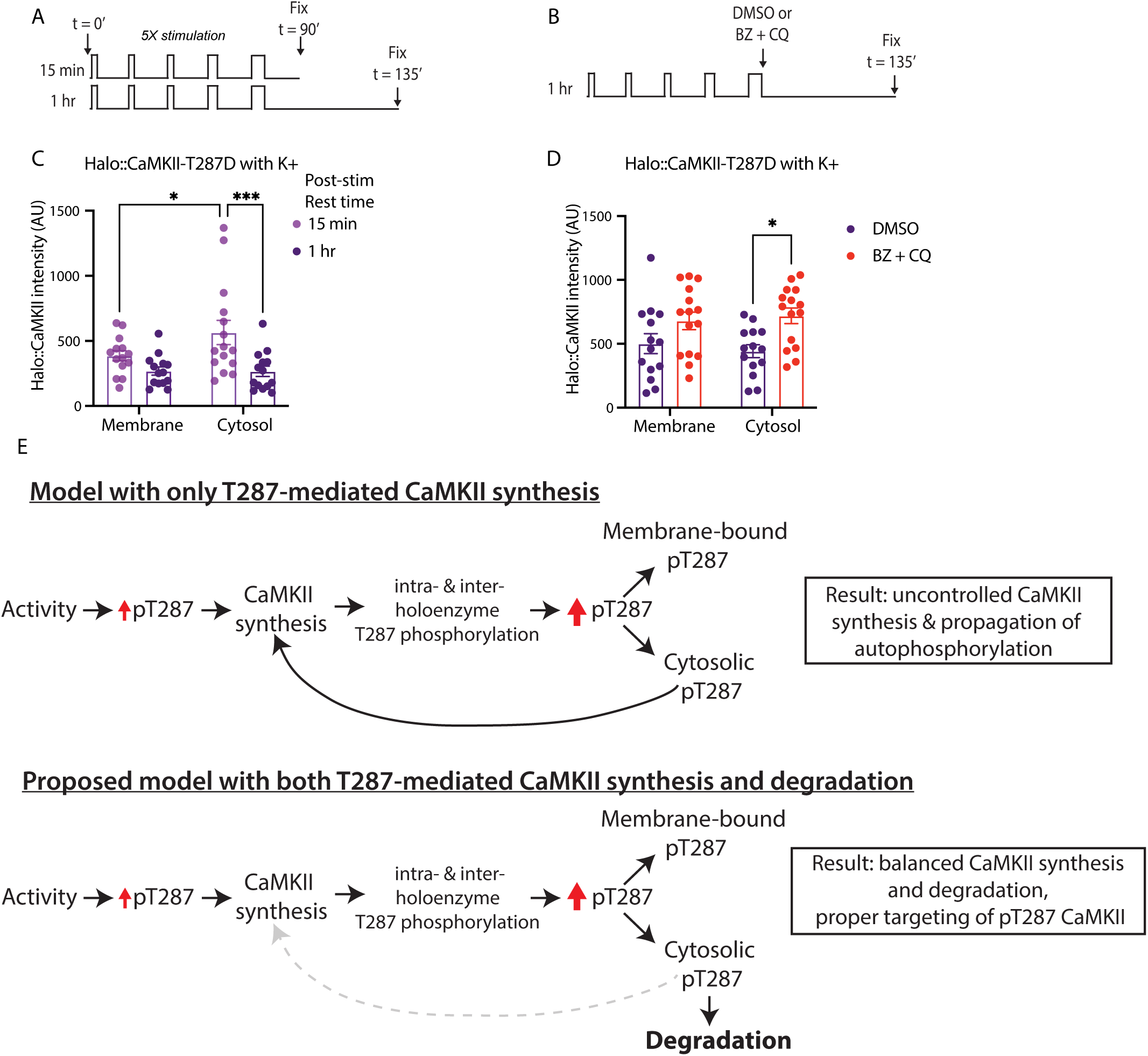
Cytosolic Halo::CaMKII^T287D^ is degraded after stimulation. (A) Halo::CaMKII^T287D^ larvae were dissected, stimulated with 5x K+ and then allowed to rest in normal HL3.1 for 15 min or 1 hr. (C) Quantification of Halo staining in the membrane or cytosol regions. n = 10 for 15 min, n = 8 for 1 hr. (B) Larvae were stimulated with 5x K+, then DMSO or Bortezomib and Chloroquine were applied in normal HL3.1 for 1 hr. (D) Quantification of Halo staining in the membrane or cytosol regions. n = 14 for DMSO, n = 15 for BZ +CQ. Groups were compared with a 2-way ANOVA with Tukey’s multiple comparisons test. Data are reported as mean ± SEM and all P values reported as follows: n.s. = not significant, * p<0.05, ** p< 0.005, *** p<0.001, **** p <0.0001. (E) Model for T287-dependent homeostatic feedback loop controlling CaMKII synthesis and degradation. Top panel depicts a model with only autophosphorylation-driven CaMKII synthesis, leading to aberrant CaMKII abundance and signaling. Bottom panel depicts our proposed model, in which T287 drives CaMKII synthesis, and is balanced by T287-dependent degradation of excess cytosolic CaMKII, leading to a balance between synthesis and degradation and properly localized CaMKII.

We hypothesized that cytosolic Halo::CaMKII^T287D^ may be more vulnerable to degradation if membrane-bound protein was engaging in protein-protein interactions which occlude the domains of activated CaMKII that stimulate degradation (32). To test this, we inhibited proteasome- and autophagy-mediated degradation during the 1 h post-stimulation rest by applying the inhibitors bortezomib (BZ) and chloroquine (CQ) (Fig. 8B). We observed a statistically significant increase in cytosolic Halo::CaMKII^T287D^ (Fig. 8D), indicating that activated CaMKII is rapidly degraded during the period of decreased activity following a period of repetitive stimulation. This implies that the steady-state differences we see between wild type and T287D CaMKII are due to cytosolic degradation of the activated form of the kinase.

Our results support a model of activity-dependent CaMKII regulation that is mediated by an autophosphorylation-dependent negative feedback loop (Fig. 8E): Neuronal activity induces CaMKII autophosphorylation, which stimulates synthesis of more protein. This new CaMKII protein can be autophosphorylated through intra- and inter-holoenzyme interactions (65, 66), or subunit exchange (67). The autophosphorylated CaMKII either associates with binding partners at the potentiated synapse or remains in the cytosolic pool. In the absence of activation-dependent degradation, this newly phosphorylated CaMKII would further promote Pi3K/Akt/mTor signaling, creating a positive feedback loop that drives continued protein synthesis and autophosphorylation (Fig. 8E, top). Instead, we propose that this unbound, excess phosphorylated protein is targeted for degradation in the cytosol (Fig. 8E, bottom). The T287-dependent control of both synthesis and degradation provides a kinase-centered mechanism for enabling precise control over CaMKII abundance and autonomous activity after stimulation.

### Membrane-localized activated CaMKII drives plasticity of spontaneous release

While the regulation of the balance of synthesis and degradation by pT287 CaMKII is important for maintaining adequate potential for future plasticity, the mechanism by which new protein synthesis and activation generate functional change is not clear. Spaced stimulation has been shown to increase miniature excitatory junctional potentials (mEJPs) in a protein synthesis-dependent manner (44) which requires the translation-promoting distal 3’UTR of CaMKII (42). Because autophosphorylation and redistribution of CaMKII have been shown to regulate neuronal function, including spontaneous release (68, 69) we asked whether T287 phosphorylation was also the functional effector for regulation of mEJP rates in our system. We utilized intracellular current clamp to record the rate of mEJPs, which indicates the spontaneous release rate at the NMJ and can change with strong spaced stimulation (44). Previous work in our lab had shown there was a depression of basal rate and a block in plasticity of mEJPs in a *CaMKII* null mutant (42). Comparison of the rates of spontaneous release between animals with presynaptically expressed Halo::CaMKII wild type and Halo:: CaMKII^T287D^ showed that the animals with activated CaMKII had a significantly higher rate (Fig. 9A-B), without differences in resting membrane potential (-64.9 ± 1.9 vs. -66.5 ± 1.8 mV, P > 0.9, n = 7 and 8). The enhanced mini frequency in T287D NMJs is occurring in a context in which there are overall lower levels of CaMKII. The fact that it is sufficient to drive an increase in mEJP rate leads us to hypothesize that it may be in part explained by relocation of autonomous CaMKII proximal to the membrane release machinery. The nature of the binding partner(s) or whether they are altered or increased by activity is unknown.

**Figure 9.**
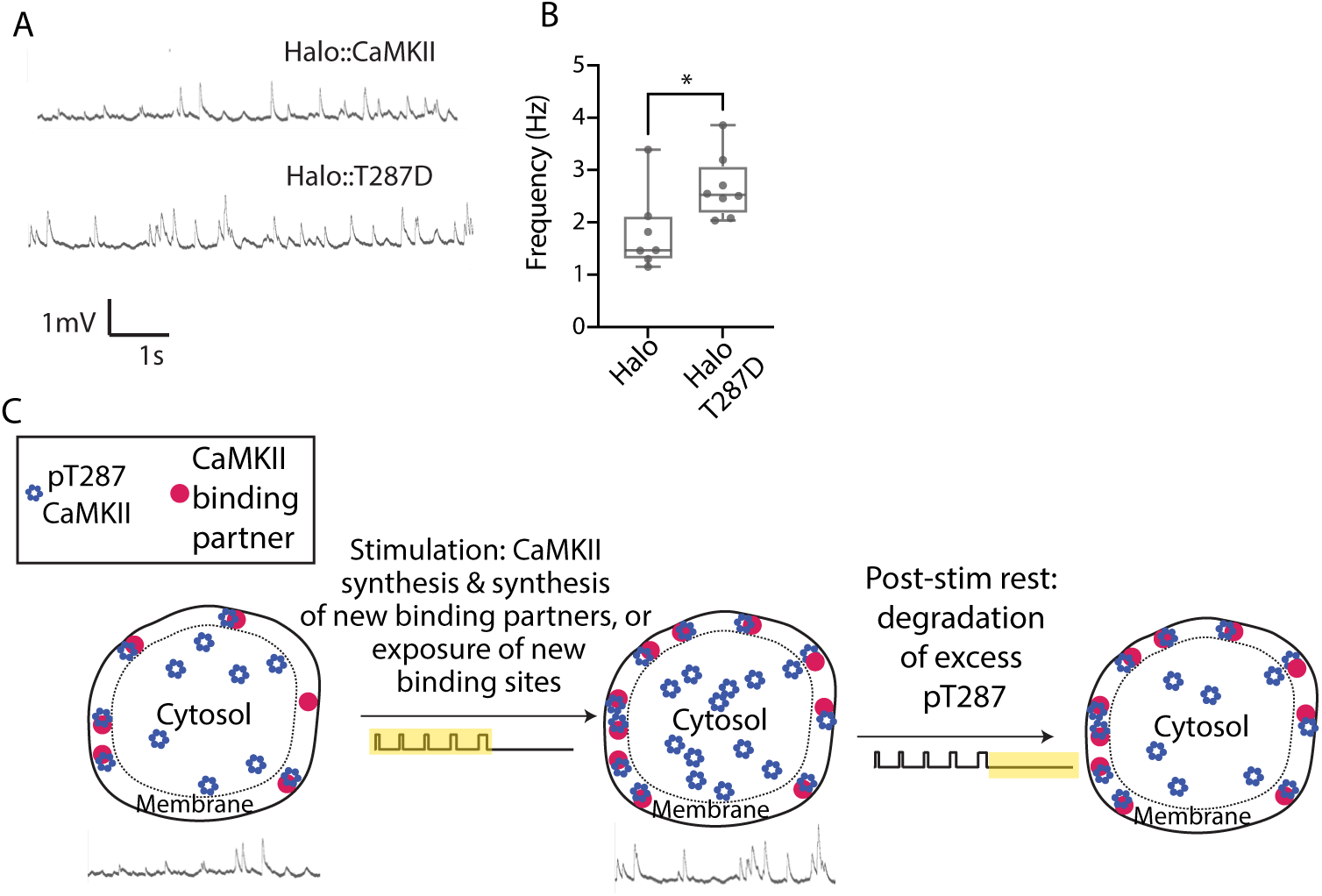
Halo::CaMKII^T287D^ causes an increase in mEJP rate in unstimulated boutons. (A) Example traces from muscle 6 of animals expressing presynaptic *Halo::CaMKII* (top) or *Halo::CaMKII^T287D^*. (B) mEJP frequency in same genotypes; Groups were compared with Welch’s t-test, n = 7 and 8 respectively. (C) Proposed model for the enhancement of mEJP frequency through membrane-localized autophosphorylated CaMKII. Activity increases both autophosphorylation and synthesis of CaMKII in the cytosol, along with the synthesis or availability of membrane binding sites. During a period of reduced activity after stimulation, excess autophosphorylated CaMKII is degraded in the cytosol. Membrane-associated CaMKII is protected from degradation due to its binding partners and contributes to enhanced mEJP frequency.

We propose that one function of activity-dependent CaMKII synthesis is to increase the pool of activated CaMKII that is available for recruitment to the synaptic membrane. These activated holoenzymes may associate with synaptic binding partners that are either newly synthesized during stimulation, or binding partners that undergo conformational changes that expose CaMKII-binding sites. During post-stimulation periods of rest, unbound activated CaMKII is degraded, restoring homeostasis of the synaptic proteome (Fig. 9C).

## Discussion

Our findings provide the first direct evidence that CaMKII is locally synthesized in the presynaptic terminal of the *Drosophila* NMJ in response to activity. Previous studies using translational reporters with an incomplete *CaMKII* 3’UTR had concluded that axonal CaMKII accumulation was not due to local synthesis (21, 24). Using endogenous locus-based tools and axotomy, we show that synthesis occurs within the presynaptic terminal and is regulated by the distal *CaMKII* 3’UTR and PI3K/Akt/mTor signaling. Additionally, we demonstrate that newly-synthesized CaMKII localizes to a different subcellular region relative to pre-existing protein, supporting the idea that the new protein may have a distinct function. We also identify multiple roles for CaMKII autophosphorylation in regulating mEJP frequency along with the synthesis, degradation and redistribution of CaMKII. These results advance our understanding of how the presynaptic cell regulates activity-dependent synthesis of protein, and how local translation of CaMKII can contribute to synaptic plasticity.

Previous work has established that *CaMKII* mRNA is present in synaptic compartments and that its local translation supports plasticity in both vertebrate dendrites (22, 38) and *Drosophila* antennal lobe and mushroom bodies, regions that are involved in memory and learning (21, 43). However, little was known about the mechanism of activity-dependent presynaptic CaMKII synthesis. Our results show that at the NMJ, presynaptic CaMKII protein levels increase in response to spaced depolarization, even when axons are severed from the somatic regions. Most of our data have utilized an acute high potassium stimulation protocol to examine activity-dependent CaMKII synthesis; however, our results indicate that the levels of CaMKII are bidirectionally influenced by chronic activity patterns at the NMJ, and that both steady-state and activity-dependent synthesis are dependent on the UTR. These results indicate that the processes that underlie activity-dependent and steady-state synthesis of presynaptic CaMKII may be similar.

Activity-dependent local translation requires that the mRNA localizes at the synapse and that there is stimulation of translation in response to activity. The 3’UTRs of mRNAs have been shown to contain sequences that are capable of regulating both of these functions by binding to RNA-binding proteins (70). Our data indicate that the long but not the short isoform of the *CaMKII* 3’UTR is involved in activity-dependent synthesis. This parallels previous evidence from our lab that determined binding of RNA binding protein Mub to a specific sequence on the 3’UTR to be crucial for steady-state CaMKII synthesis in the adult mushroom body (43). These studies along with others highlight the critical role of the 3’UTR in synaptic CaMKII translation.

An important finding of this study is that CaMKII phosphorylation at T287 affects presynaptic structure and function in multiple ways. The first is by regulating the mechanism of its own synthesis. While T287 phosphorylation has been recognized as a long-lasting molecular switch regulating learning and memory (71), our data suggest that T287 phosphorylation has an additional role in enabling CaMKII to stimulate its own synthesis through activation of the PI3K/Akt/mTor pathway. In *Drosophila* and mammalian neurons, PI3K signaling has been shown to bidirectionally regulate synapse growth (56, 72, 73) and synaptic plasticity (74, 75). Since CaMKII is an important signaling and scaffolding molecule, T287-induced stimulation of CaMKII synthesis via PI3K signaling may aid in supporting long-lasting synaptic growth following sustained neuronal activation. Additionally, because mTOR activates translation factors that broadly stimulate protein synthesis (76, 77), CaMKII T287 phosphorylation may likewise enhance the production of other key synaptic restructuring proteins that can work independently of or in conjunction with CaMKII. While spaced neuronal stimulation and PI3K/Akt/mTor signaling have separately been recognized as promoting protein synthesis and synaptic growth, our data provide a mechanism for how calcium-induced activation of CaMKII and the PI3K/mTor pathway may collaborate to lead to long-lasting protein synthesis-dependent plasticity.

Is activation of the kinase the only driver of long-term structural plasticity? Both old and new CaMKII could become activated with strong stimulation. Our data suggest that there is a more subtle process coincident with this activation and that the cell utilizes multiple pools of CaMKII protein that are spatially segregated due to their history and different phosphorylation states and/or interacting partners. This would allow for the cell to react precisely to specific stimuli. Our findings demonstrate that CaMKII synthesized following spaced stimulation constitutes an overlapping yet distinct pool from pre-existing CaMKII. These pools are likely to be directed at distinct subcellular compartments and confer specialized functions. A quantitative proteomics study of rat forebrain (13) identified a set of CaMKII interacting proteins present at substoichiometric quantities relative to CaMKII, consistent with the existence of distinct CaMKII protein pools. Here we show that these pools are spatially separated, thereby creating discrete microenvironments in which CaMKII can interact with specific sets of synaptic proteins.

The second way that T287 phosphorylation affects the synapse is likely structural. The role of T287 phosphorylation in translocation has been most convincingly demonstrated at the vertebrate postsynaptic density (8, 78). On the presynaptic side in mammals, neuronal activity (18, 79) can lead to CaMKII translocation to vesicles or presynaptic active zones, however it was not clear how T287 phosphorylation was involved. The story was also not so clear in *Drosophila.* In Shakiryanova et al. 2011, the authors showed that blocking T287 phosphorylation of a CaMKII-based sensor protein did not affect its translocation in response to a single burst of activity. In contrast, we demonstrate that expression of a phosphomimetic T287D mutant from the endogenous locus affects the distribution of CaMKII, localizing it close to the synaptic membrane where active zones are positioned. One possible explanation for these divergent findings is that the sensor used in the previous study has been shown to coassemble with endogenous CaMKII (80) which could become phosphorylated and translocate the sensor/kinase complex. We believe that it is likely that the membranous localization of T287D CaMKII in our experiments is due to kinase binding to one or more activation-specific partners at the synapse (81).

The third effect of presynaptic expression of T287D CaMKII was functional - it increased spontaneous release as measured by the rate of miniature excitatory junctional potentials (mEJPs). Spaced stimulation, either with patterned electrical stimulation or high K+ washes, has been shown to increase mEJP rate in a protein synthesis-dependent manner (44).

Previous work from our lab had shown that *CaMKII* mRNA with an intact distal UTR was required for this plastic change; CaMKII null mutants or null mutants rescued with only the short 3’UTR sequence had suppressed basal mEJP rate and could not increase the rate after stimulation (42). In the current study we show that the distal UTR is a requirement for both achieving normal basal CaMKII levels and for supporting activity-dependent synthesis, and this might be construed to mean that high levels of CaMKII protein are required to support spontaneous release. Our finding that expression of an activated T287D CaMKII from the endogenous locus is sufficient to increase mEJPs contradicts this supposition since T287D CaMKII is present at lower levels than wild type, likely due to an increased degradation rate (32). These data suggest that it is some combination of sustained CaMKII activity and membrane localization which is the key step in regulation of the mEJP rate, not the total amount of CaMKII. But because protein synthesis inhibitors can block mEJP plasticity it is clear that new protein is required, making it tempting to speculate that the presence of newly-synthesized CaMKII and/or CaMKII binding partners drive a subcellular reorganization that is critical for mEJP plasticity.

Finally, we outlined a homeostatic role for T287 autophosphorylation in CaMKII synthesis and degradation. Activation-dependent spread of autophosphorylation has been hypothesized to be a factor in the persistence of activation state despite protein turnover (66, 67, 82). However, T287-dependent synthesis combined with propagation of activated subunits may activate a positive feedback loop that saturates the cell with autophosphorylated kinase, causing promiscuous signaling, mistargeting of CaMKII protein, and/or disruption of plasticity (83, 84). While T287 dephosphorylation by protein phosphatases (85) likely occurs concurrently with synthesis and degradation, CaMKII subunits far outnumber phosphatases (86) and may be unable to keep up with the proliferation of activated subunits that may occur during strong stimulation. We propose that the combined degradation and synthesis of autophosphorylated CaMKII allows the cell to finely tune CaMKII quantity in response to the level of stimulation. Combined synthesis and degradation of CaMKII would likely aid in proper targeting of the protein. We demonstrate that the cytosolic pool of autophosphorylated CaMKII is preferentially degraded over the membrane pool. This is possibly due to activated CaMKII binding to membrane proteins, which likely prevent the degradation machinery from accessing the less stable regulatory and kinase domains of CaMKII (32). Excess subunits, which may contribute to aberrant excitability or cellular signaling, remain near the site of synthesis in the cytosol and are degraded.

CaMKII is an abundant synaptic protein, making up 1-2% of total brain protein (4). Why the synapse would synthesize more of an already abundant protein following activity remains a mystery. Our data support a model in which activity-induced CaMKII autophosphorylation regulates its synthesis, localization, and degradation, ultimately promoting the accumulation of activated CaMKII near synaptic release sites, resulting in altered synaptic function. These results provide a foundation for future studies investigating the specific functional roles of membrane-associated CaMKII and its interactors, with broader implications for understanding how activity-dependent protein synthesis contributes to long-lasting synaptic plasticity.

## Materials and Methods

### Fly cultivation

All fly lines were grown on dextrose/cornmeal/yeast food at 25°C and checked for mites and contamination regularly.

### Generation of fly lines

*CaMKII^Udel^, CaMKII^UShort^* and wild type *CaMKII^ULong^* flies were previously generated as described in Chen et al. 2022. To conditionally knockout the *CaMKII* UTR in presynaptic cells, *CaMKII^FRT−3’UTR−FRT^* flies containing two Frt site flanking the 3’UTR (also created in Chen et al. 2022) were crossed with C164-Gal4 and UAS-Flp to drive the removal of the UTR. For controls, C164-Gal4 or UAS-Flp were crossed with *CaMKII^FRT−3’UTR−FRT^* flies.

The Frt-stop-Frt-EGFP-CaMKII plasmid was created by assembling an Frt-flanked stop site, EGFP tag, and the CaMKII sequence, between two attp sites into a pBS-KS-attB2 plasmid. The Frt-stop-Frt-Halo-CaMKII plasmid was generated from the Frt-stop-Frt-EGFP-CaMKII plasmid by excision of the EGFP tag and replacement with a Halo Tag cloned from the plasmid UAS-Halo7::CAAX (gift from Gregory Jefferis - Addgene # 87645). The T287D mutation was then inserted into the plasmid with site-directed mutagenesis. The sequences of the plasmids were confirmed by whole-plasmid sequencing, and then injected into *CaMKII^coding−3P3−RFP^* flies (Chen et al. 2022) by Rainbow Transgenic Flies. The insertion of the Halo::CaMKII cassette into the genome was confirmed by the lack of RFP expression. After selection, the correct insertion sequence in candidate fly lines was confirmed with Sanger sequencing with primers that bind outside of the integration sites.

*CaMKII^T287T^, CaMKII^T287A^*, and *CaMKII^T287A,T306,7A^* flies were created by inserting either the wildtype CaMKII sequence (T287T), or mutated sequence (T287A and T287A, T306/7AA) into *CaMKII^coding−3P3−RFP^*flies. Correct insertion sequence in candidate fly lines was confirmed with Sanger sequencing with primers that bind outside of the integration sites. *eag^1^sh^120^*and *para^ts1^* flies were a gift from Troy Littleton (MIT).

### Spaced depolarization/high potassium stimulation

Crosses consisting of four female and four female flies were transferred to a new culture tube every two days to ensure that the density of larvae were consistently sparse between experiments. Once wandering third instar larvae appeared, they were collected, and the sex of the larvae determined. Females were used because they exhibit more robust plasticity than males (87).

Normal and high K+ HL3.1 solutions were prepared the same day of the experiment. The composition of the normal and high potassium HL3.1 was modeled on (44) and optimized for the induction of protein synthesis by using a reduced magnesium concentration to allow for greater excitability. (Molarity in mM as follows - Normal HL3.1: 70 NaCl, 5 KCl, 0.1 CaCl2, 10 MgCl2, 10 NaHCO3, 5 trehalose, 107 sucrose, 5 HEPES-NaOH, pH 7.2; and High K+ HL3.1: 40 NaCl, 85 KCl, 1 CaCl2, 10 MgCl2, 10 NaHCO3, 5 trehalose, 5 sucrose, 5 HEPES-NaOH, pH 7.2). Solutions were checked for correct osmolarity (+/- 5 mOsm) with an osmometer.

Larvae were dissected in ice cold normal HL3.1, leaving the central nervous system and axons intact. They were stretched to only 60% of their maximal stretching capacity to allow for contractions during stimulation. After dissection, they were gently washed with normal HL3.1 to remove excess debris. The spaced depolarization was performed as described in Ataman et al. 2008: the normal HL3.1 was replaced with high K+ HL3.1 for pulses of 2 min, 2 min, 2 min, 4 min, 6 min, with rest in normal HL3.1 in between the pulses and at the end for 15 min. For the normal condition, the high K+ washes were omitted and replaced with normal HL3.1 washes. For single color HaloTag experiments (Figs. 8A-D), the HaloTag ligand (200 nM) was applied before stimulation for 15 min, washed 3x in normal HL3.1 The ligand was reapplied at the penultimate rest to ensure complete staining. For pharmacology experiments, the drug was included in all normal and high K+ incubations after dissection, except for experiments with Bortezomib and Chloroquine (8D), which were only included in the final rest period.

Concentrations and sources of drugs in solution were as follows: Cycloheximide (1 mM, ThermoFisher), EGTA (0.5 mM, Sigma), Anisomycin (40 µM, Sigma), Wortmannin (100 nM, Sigma), Rapamycin (100 nM, ThermoFisher), Bortezomib (100 nM, MedChemExpress), Chloroquine (50 µM, Sigma). At the end of the stimulation protocol, the larvae were fully stretched to allow for examination of NMJ morphology and protein levels, and then fixed in 4% paraformaldehyde in normal HL3.1 for 25 min. The fixative was removed by washing in PBX (PBS pH 7.4 with 0.1% Triton X-100) 3 times for 10 min each.

### Immunostaining

Fixed larval fillets were stained in primary antibody in PBX with 10% normal goat serum overnight at 4°C while gently shaking. Excess antibody was removed with 3 washes in PBX for 10 min each. Secondary antibody was applied in PBX for 2 hours at room temperature, and then washed 3 times for 10 min each. The fillets were mounted in SlowFade Diamond AntiFade Mountant (ThermoFisher), sealed with clear nail polish and stored at 4°C until imaging. Slides were imaged within 1-2 days of mounting to preserve fluorescent signal. Antibody dilutions are as follows: Ms anti-CaMKII (#18, CosmoBio) 1:1000; Ms anti-phospho CaMKII (22B1, ThermoFisher) 1:100; Rb anti-HaloTag (G9281, Promega) 1:500; Rb anti-Phospho Akt (4060T, CST) 1:500; Rb anti-GFP (A-11122, ThermoFisher) 1:1000; Ms anti DLG (4F3, DHSB) 1:100; Rb anti-DLG 1:1000 (Boster Bio); Rb anti-HRP (Jackson Immunoresearch); Ms Anti-Brp (nc82, DHSB) 1:100; Alexa Fluor 647-conjugated anti-HRP (Jackson) 1:250; Alexa Fluor 488-conjugated anti GFP (A21311, ThermoFisher) 1:250; Goat anti-Mouse Alexa Fluor 488 (A11029, ThermoFisher) 1:250; Goat anti-Rabbit Alexa Fluor 488 (A11008, ThermoFisher) 1:250; Goat anti-Rabbit Alexa Fluor 555 (A21428, ThermoFisher) 1:250; Goat anti-Rabbit Alexa Fluor 647 (A21244, ThermoFisher) 1:250; Goat anti-Mouse Alexa Fluor 647 (A21240, ThermoFisher) 1:250.

### Electrophysiology

All electrophysiological recordings were carried out at room temperature in modified HL3.1 saline (70 mM NaCl, 5 mM KCl, 4 mM MgCl₂, 10 mM NaHCO₃, 115 mM sucrose, 5 mM trehalose, 5 mM HEPES, 1.5 mM CaCl₂, pH 7.2) (88). Following the electrical stimulation paradigm reported by Ataman et al. 2008 (44), the uncut A3 segmental nerve was gently drawn into a suction electrode. Stimulation was delivered with a Master-8 pulse generator (A.M.P.I.) driving an A-M Systems Model 2100 stimulator. Preparations received 10 Hz stimulation for 2s followed by 3s of rest, repeated continuously for 5 min (40% duty cycle). After a 15-min rest, this 5 min stimulation block was repeated four additional times. After the final block, tissues were fixed in 4% paraformaldehyde in HL3.1 for 20 min at room temperature, washed in PBX and processed for immunostaining.

Miniature excitatory junctional potentials (mEJPs) were recorded using an Axon Instruments MultiClamp 700A amplifier. Sharp-electrode recordings of neuromuscular junctions were obtained from muscle 6 in abdominal segment A3 using glass microelectrodes with a resistance of 12–20 MΩ. Recordings were included for analysis only if the resting membrane potential was more hyperpolarized than −59 mV and the input resistance exceeded 4 MΩ. Spontaneous mEJP frequency was quantified by measuring the time required to acquire the first 200 events. Events were detected using a custom MATLAB script based on code from the Marder lab (https://github.com/marderlab/Spike_analysis).

### HaloTag Pulse Chase

Larvae were dissected and then rinsed with normal HL3.1. The pulse (“old”) HaloTag ligand (Janelia Fluor 646, Promega #GA1120) (133 nM) was applied for 15 minutes, followed by three quick rinses with HL3. After a 5-minute incubation in HL3.1, the chase (“new”) HaloTag (Janelia Fluor 549, Promega # GA1110) (500 nM) was applied for 15 minutes. After three quick rinses, and one 15 min rinse in normal HL3.1, high potassium stimulation was performed as described in the previous section. For the penultimate rest, the chase HaloTag was reapplied (250 nM) to ensure complete labeling of all CaMKII. Larvae were stretched, fixed in 4% paraformaldehyde in normal HL3.1 for 25 min. After washing in PBX 3 times for 10 min each, they were processed for immunohistochemistry.

### HCR-FISH

Reagents and probes were purchased from Molecular Instruments and RNase-free water was used for all solutions. Larvae were dissected in cold HL3.1 as previously described. They were fixed in 4% PFA for 35 minutes and then washed in PBST (PBS pH 7.4 with 0.5% Tween-20) two times for 20 minutes each. After one wash in PBS, they were incubated in 10 µg/ml Protease K in PBS for 10 min at 37°C. The protease was removed with three washes in PBS and the samples were then processed according to the Molecular Instruments kit protocol.

### Microscopy

For all experiments, muscle 6/7 NMJs from segments A3 and A4 were imaged for each larvae. For experiments with presynaptically-or postsynaptically-expressed EGFP::CaMKII (Figs. 1, 2A-H, 3A-F), Z-stacks of NMJs were acquired with a Leica SP5 confocal microscope using the 20x objective and 4x zoom and the 488 or 633 laser lines. Fig. 2K-L was acquired on Zeiss LSM880 without AiryScan using the 63x/1.4NA objective.

For analysis of CaMKII antibody or HaloTag staining and HCR-FISH (Figs. 3H-I, 4–8, in which high resolution of the NMJ at the bouton level was required, Z-stacks of larval NMJs were acquired with a Zeiss LSM880 Airyscan fast confocal system. The 63x/1.4NA objective and the appropriate laser line was used (488, 561, or 633 nm). After acquisition, the Z-stacks were processed using standard Airyscan processing.

### Image analysis

To quantify the fluorescent intensity of larvae expressing Halo-or EGFP-tagged CaMKII (Figs. 1, 2, 3A-F), files were opened in Fiji. The section containing the bouton center was identified by Hrp or Dlg staining. The image was processed with a Gaussian blur filter and thresholded in the Hrp or Dlg channel using the Otsu method. Regions of interest (ROIs) were then identified with the “Analyze Particles” function, and the mean intensity measured. Each ROI was manually verified to ensure it corresponded to the presynaptic region.

To quantify the levels of immunostained CaMKII, pT287 or pAkt in *Canton S* larvae or total HaloTag levels in Halo::CaMKII larvae (Figs. 3H-I, 4, 5C-D, 6A-D), each bouton was manually outlined and added to the ROI manager. The multiple ROIs were combined into one ROI using the “OR (Combine)” function, and then the mean intensity was measured.

To compare the staining intensity of the membrane and cytosol regions (Figs. 6F, 7–8), an ROI was drawn around one bouton. The “Scale” function was used to create another ROI that is 0.7 the area of the whole bouton, which was considered the cytosolic region. The membrane ROI was created by subtracting the cytosolic region from the whole bouton using the “XOR” function. The mean intensities could then be calculated using “Multi Measure”.

For Fig. 5G-J, the intensity throughout the Z axis was determined by zooming in on a single terminal bouton, and outlining the ROI and adding to the ROI manager. Using the function “Plot Z-axis Profile” the mean intensity within the ROI of each Z slice was determined for each channel. The Z-axis profiles of each bouton were aligned by determining the central Z slice of the bouton, which corresponded to the maximum diameter of the bouton in the X-Y plane in the Brp channel.

For HCR-FISH experiments (Fig. 2I-J), mRNA puncta density (per µm³) was quantified separately for presynaptic and postsynaptic regions. In presynaptic boutons, puncta within a 1 µm-thick cylindrical section centered on the bouton were counted, with adjacent slices examined to confirm puncta localization within the bouton rather than the subsynaptic reticulum.

For all analyses, the larvae were randomly assigned a number and the experimenter was blinded to the conditions until the analysis was completed. The data was analyzed with GraphPad Prism using the appropriate statistical test.

## Acknowledgments

We thank Dr. Andrew Stone from the Brandeis Light Microscopy Core Facility (RRID:SCR_025892) for help with imaging and Julia Birnbaum for assistance with generating figures.

## Funding Sources

This work was funded by NIH R37 NS112810 (to LCG), F31 NS134189 (to KC), T32 NS007292 (to KMDLG) and R01 NS116375 (to AAR).

---

**Supplemental figure 1. Areas under the curve (AUCs) indicate difference in quantity of new, but not old CaMKII in individual boutons.** Areas under the curve for each bouton were generated for old CaMKII (A), new CaMKII (B) or new/old CaMKII ratio (C) using GraphPad Prism with threshold = 0. Groups were compared using Mann-Whitney tests. Data are reported as mean ± SEM and all P values reported as follows: *** p<0.001, **** p <0.0001.

**Supplemental figure 2. Activity increases total CaMKII and pT287-CaMKII in the cytosol.** (A, C) Quantification of pT287 (A) or total CaMKII (C) in the membrane and cytosolic regions in normal or stimulated K+ NMJs. (B, D) Representative single boutons from experiments of anti-pT287 (B) or anti-CaMKII staining. Scale bars = 2 μm. (D). n= 15 for all groups. Groups were compared with 2-way ANOVA with Tukey’s multiple comparisons test. Data are reported as mean ± SEM and all P values reported as follows: n.s. = not significant, * p<0.05, ** p< 0.005, *** p<0.001, **** p <0.0001.

## Notes

### Competing Interest Statement

The authors have declared no competing interest.

### Summary of Updates

Several small edits were made in order to clarify the differences in analyses between figure 5 and later figures.

