## Supplementary figures and images for "Activity drives local CaMKII synthesis and subcellular localization via autophosphorylation-dependent pathways"

### Figure S1

Figure S1.

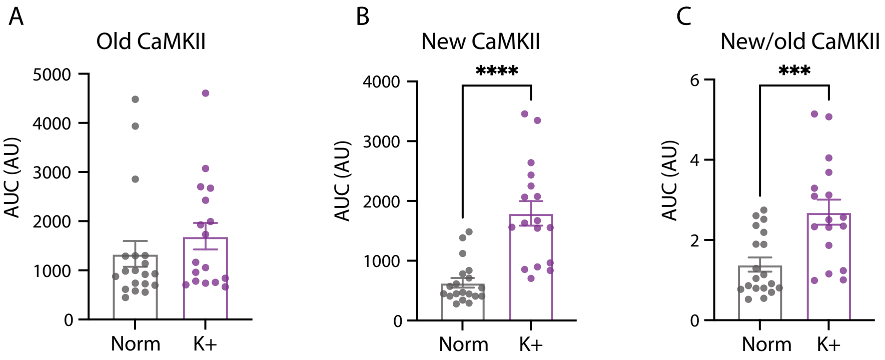
A
Old CaMKII
5000
4000
3000
2000
1000
0
AUC (AU)
Norm
K+
B
New CaMKII
****
4000
3000
2000
1000
0
AUC (AU)
Norm
K+
C
New/old CaMKII
***
6
4
2
0
AUC (AU)
Norm
K+
