## Supplementary material for "Activity drives local CaMKII synthesis and subcellular localization via autophosphorylation-dependent pathways": Figure S2

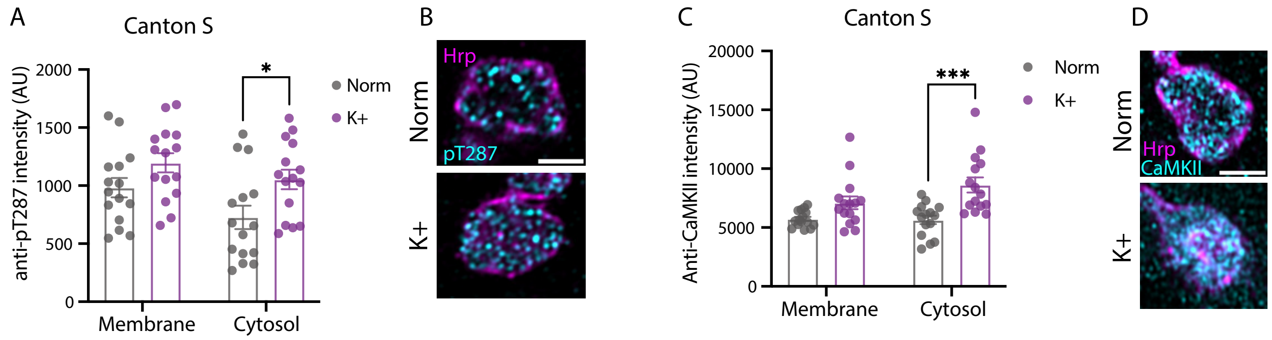
A
Canton S
anti-pT287 intensity (AU)
2000
1500
1000
500
0
*
Norm
K+
Membrane
Cytosol
B
Hrp
pT287
Norm
K+
C
Canton S
Anti-CaMKII intensity (AU)
20000
15000
10000
5000
0
***
Norm
K+
Membrane
Cytosol
D
Norm
Hrp
CaMKII
K+
